# TransGeneSelector: A Transformer-based Approach Tailored for Key Gene Mining with Small Plant Transcriptomic Datasets

**DOI:** 10.1101/2023.09.26.559592

**Authors:** Kerui Huang, Jianhong Tian, Lei Sun, Peng Xie, Shiqi Zhou, Aihua Deng, Ping Mo, Zhibo Zhou, Ming Jiang, Guiwu Li, Yun Wang, Xiaocheng Jiang

## Abstract

Gene mining, particularly from small sample sizes such as in plants, remains a challenge in life sciences. Traditional methods often omit significant genes, while deep learning techniques are hindered by small sample constraints and lack specialized gene mining approaches. This paper presents TransGeneSelector, the first deep learning method tailored for key gene mining in small transcriptomic datasets, ingeniously integrating data augmentation, sample filtering, and a Transformer-based classifier. Tested on *Arabidopsis thaliana* seeds’ germination classification using just 79 samples, it not only achieves classification performance on par with, if not superior to, Random Forest and SVM but also excels in identifying upstream regulatory genes that Random Forest might miss, and these pinpointed genes more accurately reflect the metabolic processes inherent in seed germination. TransGeneSelector’s ability to mine vital genes from limited datasets signifies its potential as the current state-of-the-art in gene mining in small sample scenarios, providing an efficient and versatile solution for this critical research area.

## Introduction

Gene mining encompasses a series of methods that unveil genes playing critical roles in specific biological processes, a task of profound importance in the field of life sciences. In plants, many vital agronomic traits, such as yield and disease resistance, are manifested as complex quantitative characters, governed by multiple genes in conjunction with environmental interactions(Fregene et al., 2001; Wang et al., 2017; Westerman et al., 2022). Enhancing these traits necessitates the preliminary identification of their underlying genetic mechanisms and key regulatory genes. In the realm of medical research, numerous diseases have genetic underpinnings, with an individual’s susceptibility often linked to variations in genes and gene expression patterns(Su et al., 2020). The pursuit of mining disease-related genes, and deciphering their roles in associated cellular processes and signaling pathways, serves to deepen our understanding of disease pathogenesis(Su et al., 2020), pinpoint disease-related biomarkers(Wang et al., 2021), and potentially pave the way for identifying novel therapeutic targets(Florez, 2017).

Gene mining often start with small-sample studies due to the limited availability of samples and the high cost of sequencing. Consequently, conventional transcriptional data mining approaches often rely on the fold change of gene expression levels and functional enrichment to identify differentially expressed genes(Mutz et al., 2013; Cao et al., 2014; Chen et al., 2020). However, this is a rudimentary method that tends to overlook many vital genes. Contrasted with traditional methods, machine learning techniques offer a more refined approach. They can effectively extract key genes related to specific phenotypes or diseases as biomarkers from high-dimensional transcriptome data, achieving high accuracy and effectiveness. This approach is currently a favored method in key gene mining(Su et al., 2020). For instance, Li et al. utilized machine learning to identify key genes from gene expression data of COVID-19 patients to assess disease severity(Li et al., 2022). Yu et al. applied machine learning and transcriptome sequencing to discern 9 SNPs in the transcriptome data of Platycodon grandiflorus, aiding in flower color identification(Yu et al., 2021). Additional works include the development of prediction tools for disease-resistant proteins in plants by Pal et al. based on support vector machines(Pal et al., 2016), and Chen et al.’s innovative machine learning method combinations to identify key genes in bovine multi-tissue transcriptome data for predicting feed efficiency(Chen et al., 2021).

Although the marriage of transcriptome sequencing and machine learning has shown promise in key gene mining, traditional machine learning algorithms often require manual feature engineering to enhance performance. This practice leads to the discarding of many genes that might seem unimportant on the surface. However, considering the complex regulatory relationships and concealed action patterns that genes in organisms typically exhibit(Karlebach and Shamir, 2008; Crombach et al., 2012), these disregarded genes might possess key value. The limitations inherent in traditional machine learning underline the importance of developing algorithms capable of mining key genes from transcriptomics data while fully capturing the intricate global interactions between genes.

Deep learning, a subset of machine learning, employs artificial neural networks (ANNs) or deep neural networks (DNNs) to model complex problems. Unlike traditional techniques, deep learning can process unstructured data and automatically learn effective features from high-dimensional data without manual feature selection, making it well-suited for mining key genes in biological processes(Wu and Chen, 2015; Sau and Balasubramanian, 2016; Suryanarayana et al., 2018; Sakellaropoulos et al., 2019; Pacal et al., 2020; Saxe et al., 2020). Among deep learning approaches, natural language processing (NLP) models like RNN(Chung et al., 2014; Shewalkar et al., 2019) and LSTM(Yu et al., 2019) are notable for their ability to capture long-distance dependencies in sequences. This characteristic can be applied to unravel the complex regulatory relationships between genes. The Transformer model stands out as a significant advancement in this field. Introduced in 2017, this model is designed for NLP tasks and relies on an attention mechanism, surpassing traditional RNNs and LSTMs in capturing long-range dependencies between sequential elements(Vaswani et al., 2017). The Transformer’s impact has been profound, revolutionizing the field of NLP(Ma et al., 2019; Yan et al., 2019; Schrimpf et al., 2021). Consequently, the application of Transformers in classifying biological processes and mining key genes offers a promising and potentially groundbreaking avenue for research.

Studies to date integrating transcriptomic gene expression data with Transformer architectures have primarily centered on single-cell sequencing, specifically for cell type classification(Chen et al., 2023; Xu et al., 2023). Examples include TOSICA, a Transformer-based model for cell type annotation, and STGRNS for inferring gene regulatory networks. Although valuable, these works have not tackled key gene mining, and with the high costs of single-cell sequencing, the majority of biological studies still rely on traditional transcriptomics sequencing. There’s a strong practical need for developing methods to mine key genes using Transformer architectures with traditional transcriptomics sequencing data. Only two studies have applied Transformers to normal RNA sequencing for cancer phenotype prediction and cancer type classification(Zhang et al., 2022; Khan and Lee, 2023), but neither focused on key gene mining.

One challenge in integrating standard transcriptomics sequencing data with deep learning lies in the limited sample size of standard transcriptomics sequencing data, which contrasts with the high sample demands of deep learning(Milicevic et al., 2018; Reyes-Nava et al., 2018). This limitation is why traditional machine learning is more commonly used, as it doesn’t require a large number of samples(Xiao et al., 2015; Rajput et al., 2023). With the growth of deep learning, data augmentation techniques such as GANs have been employed to enhance transcriptomics data(Liu et al., 2019; Shorten and Khoshgoftaar, 2019; Marouf et al., 2020). WGAN and its improved version, WGAN-GP, have shown notable improvements in sample quality and training stability(Arjovsky et al., 2017). Recently, WGAN-GP’s application to augment transcriptomics data demonstrated that artificially generated samples could enhance classification model performance. Hence, enhancing small-scale transcriptomics sequencing data with WGAN-GP, followed by the utilization of Transformer models for classifying biological processes and mining key genes, presents a feasible and promising approach.

In response to the challenges and limitations identified in traditional machine learning and deep learning applications to small transcriptomic datasets, we present TransGeneSelector. This novel approach is the first deep learning method specifically tailored for key gene mining in small transcriptomic datasets. It combines a Transformer architecture with a sample generation network based on WGAN-GP and a sample filtering network. The process starts by employing WGAN-GP to generate transcriptomic samples, followed by a filtering stage to exclude low-quality samples. Subsequently, a Transformer is utilized to classify biological processes by capturing the complex global relationships between genes. The significance of each gene is further assessed using SHAP (SHapley Additive exPlanations), a method that provides interpretative insights into individual predictions from machine learning models(Lundberg and Lee, 2017; Rodríguez-Pérez and Bajorath, 2019).

TransGeneSelector not only predicts the seed states (either dry or germinating) of *Arabidopsis thaliana* with performance comparable to the Random Forest and SVM algorithm, but it also identifies genes at higher regulatory levels that are more representative of seed germination. TransGeneSelector thus offers the following advantages:

1. Ability to analyze small sample size transcriptomic data, classify specific biological processes with high performance, and identify key genes.
2. Capability to detect vital regulatory relationships between genes, including upstream key genes that govern specific biological processes, surpassing the abilities of Random Forest.

In essence, TransGeneSelector presents a practical tool for life science researchers, particularly those in the plant domain, to mine key genes from transcriptome data, providing insights into specific biological processes.

## Result

### Overview of TransGeneSelector

TransGeneSelector is a composite deep learning method designed for classifying biological processes and identifying key genes. It integrates three distinct networks: a sample generation model using WGAN-GP, a Transformer-based classification model, and an additional fully connected classifier network (Fig. 1). The method unfolds in three main stages:

1. Fake Sample Generation (Fig.1a): The WGAN-GP model is initially trained on the training set to generate fake transcriptomic gene expression samples (fake samples I). These are combined with real samples and trained in the additional classifier to enhance distinction between fake and real samples. Subsequent filtering of fake samples II through this classifier weeds out low-quality samples, resulting in high-quality final fake samples.
2. Transformer Classification (Fig.1b): The final fake samples are mixed with real training set samples to train a simplified Transformer classification model, retaining only the encoder part. A full connected (FC) layer reduces each sample’s gene expression level to 72 dimensions, preserving global expression information. These vectors are input to an 8-layer stacked Attention head in the Transformer encoder, post Positional Encoding. The first token’s vector of the output is used to assess the classification performance on the validation set real samples.
3. SHAP Method for Key Gene Mining: The SHAP (SHapley Additive exPlanations) method evaluates each gene’s influence on the trained Transformer classification model. Based on Shapley values, the genes exerting the most significant impact on the classification are identified as key genes for specific biological processes.

**Fig. 1.**
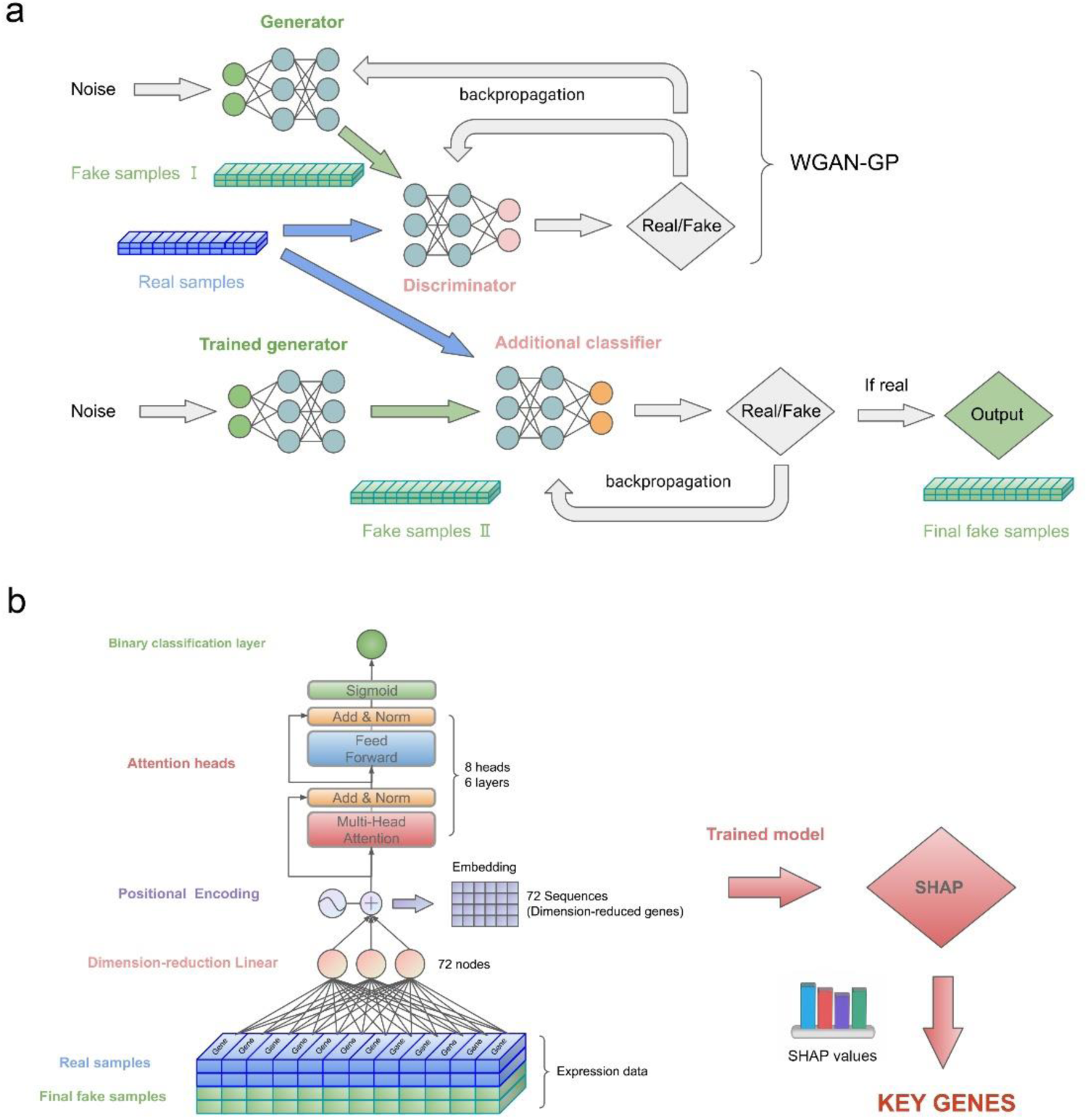
The overall frame and workflow of TransGeneSelector. **a** Network for Generating Synthetic Samples: Utilizing the WGAN-GP and additional classifier sample filtering networks, the WGAN-GP model is trained with real sample data to create synthetic gene expression samples, referred to as fake samples I. These are then amalgamated with the training set of real samples and passed through the additional classifier to enhance its discrimination capabilities between fake and real samples. Following the training of the additional classifier, fake samples II generated by WGAN-GP are processed to filter out substandard samples, resulting in high-quality, final fake samples. **b** Transformer Classification Model and SHAP Method for Key Gene Mining: The final fake samples are blended with the training set of real samples and used to train a specially simplified Transformer classification model suitable for small-sample classification tasks. The Sigmoid function is applied to the first token of the output, serving as an activation mechanism. Ultimately, the SHAP (SHapley Additive exPlanations) method is employed to evaluate the influence of each gene on the trained Transformer classification model. This results in the identification of genes with the most substantial impact on classification outcomes, facilitating the mining of key genes pertinent to a specific biological process.

### TransGeneSelector achieves high classification performance in small samples

Initially, we trained TransGeneSelector using a gene expression dataset from *A. thaliana* dry seeds (negative samples, 36 samples) and germinating seeds (positive samples, 43 samples), comprising 79 samples in total. The training of the WGAN-GP module exhibited stability and good performance after 3,800 training epochs, with the Generator and Discriminator’s Loss reaching a convergence state (Fig. 2a). Furthermore, after training for 3,800 epochs using these parameters, the Fréchet Inception Score value reached its lowest level (Fig. 2b), and the distribution of generated samples closely matched the distribution of real samples (Fig. 2c). These parameters were then utilized for subsequent network training.

**Fig. 2.**
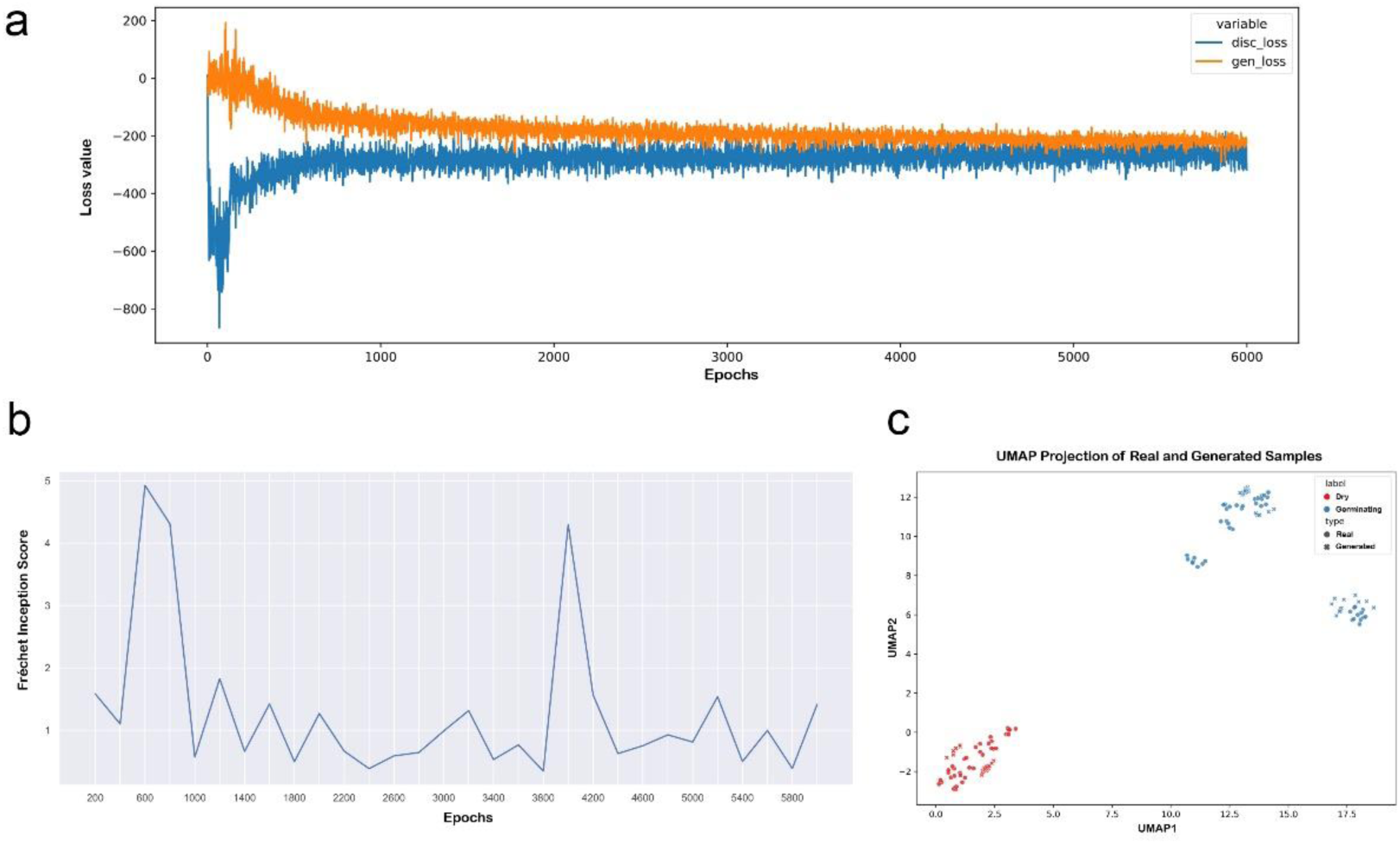
Training process and sample quality evaluation of WGAN-GP. **a** Graphical representation of the relationship between the number of training epochs and the loss values for WGAN-GP. The discriminator’s loss (disc_loss) and the generator’s loss (gen_loss) are plotted separately, reflecting the dynamics of the training process. **b** A plot showcasing the relationship between the number of training epochs and the Fréchet Inception Score for WGAN-GP, providing a measure of generated sample quality. **c** UMAP (Uniform Manifold Approximation and Projection) visualization of both the generated and real samples. Colors are used to differentiate between seed states (dry seeds or germinating seeds), while distinct shapes indicate whether samples are generated or real.

Upon fully training the WGAN-GP, we employed the real samples from the training set and an equal number of fake samples generated by WGAN-GP to train an additional classifier network for 50 epochs. This helped the network thoroughly learn the distinct attributes of both genuine and fake samples. Upon completion of this training, we set a threshold of 0.1 and selected varying numbers of high-quality fake samples generated by WGAN-GP. These were combined with real samples and inputted into TransGeneSelector’s Transformer network for training. The validation set was then applied to assess accuracy, precision, recall, and the F1 index, utilizing 5-fold cross-validation for each sample quantity.

The results (Fig. 3a) revealed that with the increase in fake samples, all cross-validation indicators initially rise, and then oscillate between increasing and decreasing. Regardless of the number of samples generated or the inclusion of an additional classifier, the model’s accuracy, precision, and F1 index consistently outperformed those of the model without any generated samples, which achieved metrics of only 0.5558, 0.4518, and 0.5714, respectively. Notably, the model that included an additional classifier and 2200 fake samples achieved the highest values: 0.9875 for accuracy, 1.000 for precision, and 0.9818 for recall and 0.9904 for F1 index. This indicates that the WGAN-GP network boosts the Transformer classification model’s performance, with an optimal 2200 fake samples. Moreover, models using an additional classifier generally outperformed those without, reducing the standard deviation of most indicators and thereby enhancing the model’s stability and reliability.

**Fig. 3.**
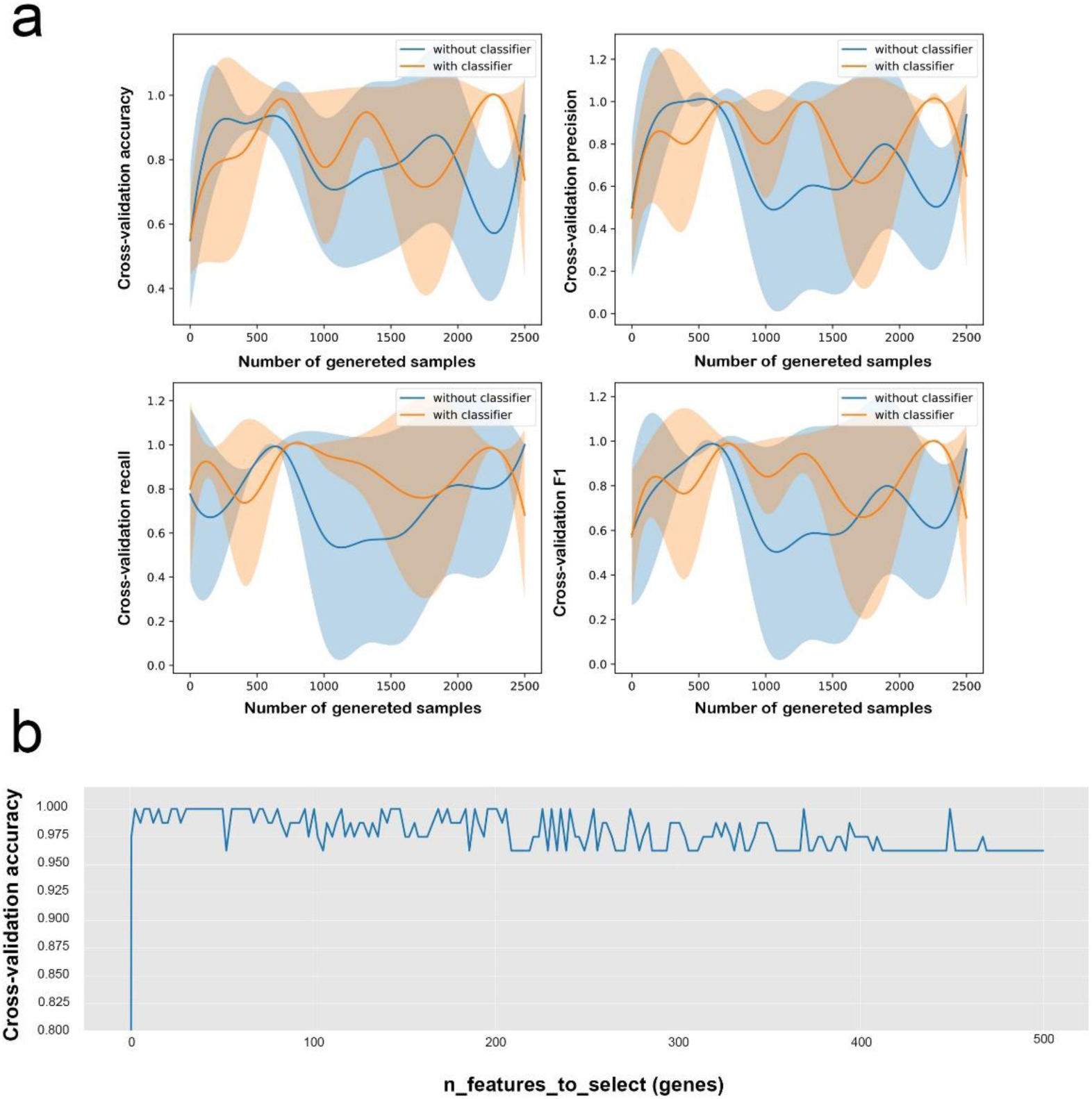
Comparison of classification performance of TransGeneSelector and Random Forest. **a** TransGeneSelector Performance Across Different Numbers of Generated Samples: This panel illustrates the classification performance of TransGeneSelector with varying numbers of synthetic samples. The blue line represents the cross-validation performance of the model without additional classifier filtering, and the orange line signifies the cross-validation performance with additional classifier filtering. The shaded regions around each line indicate the corresponding error bands, providing a visual representation of the uncertainty associated with each measurement. **b** Random Forest Classifier Performance with Varied Gene Selection: This part demonstrates the classification performance of the Random Forest classifier for different numbers of genes selected through the wrapper method. The parameter ‘n_features_to_select’ within the RandomForestClassifier module is set uniformly spaced between 1 and 500, with a specific value chosen as 200, allowing for a detailed exploration of the effect of gene selection on classifier performance.

We also employed Random Forest to classify the same dataset into germinating seeds and dry seeds. Using the Wrapper method, we selected 200 uniformly spaced parameters (number of genes) between 1 and 500 for feature engineering and classification evaluation in Random Forest, evaluating the model’s performance through cross-validation. The cross-validation results (Fig. 3b) showed that the accuracy of Random Forest could reach 1.000 with certain gene features, demonstrating the efficacy of traditional machine learning algorithms on small sample datasets.

We further utilized the test set for the definitive evaluation of our model’s performance (Table 1). For TransGeneSelector, we opted for both an unoptimized model (without the implementation of ‘early stop’ optimization) and the aforementioned model that exhibited the best cross-validation results when trained with 2,200 imputed samples. As for Random Forest, we used models trained with features that all achieved a 100% cross-validation accuracy, specifically those with 8, 11, 41, 51, 128, and 449 features, under the Wrap feature engineering method. Additionally, for a broader comparison, we incorporated a nonlinear SVM, ensuring it was trained using the same features selected through Random Forest’s feature engineering process. The assessment results indicated that TransGeneSelector had notable advantages in both accuracy and precision, each recording a peak value of 0.9524. SVM models trained using 148 and 449 features exhibited a pronounced lead in accuracy, recall, and F1 score, attaining respective maximums of 0.9524, 1.0000, and 0.9545. The Random Forest models, on the other hand, stood out solely in the AUC metric, with the model based on 8 features achieving a top AUC of 0.9875. In summary, our developed TransGeneSelector displayed evident superiority in specific metrics, ensuring that its performance in smaller sample scenarios is on par with, if not superior to, conventional methods.

**Table 1.** Quantitative Assessment of Model Performance.

| Methods | Accuracy | Precision | Recall | F1 | AUC |
| --- | --- | --- | --- | --- | --- |
| TransGeneSelector<br>(unoptimized) | 0.9048 | 0.9048 | 0.9048 | 0.9048 | 0.9229 |
| TransGeneSelector | **0.9524** | **0.9524** | 0.9524 | 0.9524 | 0.9615 |
| Random_Forest<br>with 8 genes | 0.9048 | 0.8696 | 0.9524 | 0.9091 | **0.9875** |
| Random_Forest<br>with 11 genes | 0.8810 | 0.8333 | 0.9524 | 0.8889 | 0.9841 |
| Random_Forest<br>with 41 genes | 0.9286 | 0.9091 | 0.9524 | 0.9302 | 0.9671 |
| Random_Forest<br>with 51 genes | 0.9286 | 0.9091 | 0.9524 | 0.9302 | 0.9660 |
| Random_Forest<br>with 128 genes | 0.9286 | 0.9091 | 0.9524 | 0.9302 | 0.9615 |
| Random_Forest<br>with 449 genes | 0.9286 | 0.9091 | 0.9524 | 0.9302 | 0.9626 |
| SVM with 8 genes | 0.8571 | 0.8000 | 0.9524 | 0.8696 | 0.9263 |
| SVM with 11 genes | 0.8571 | 0.8000 | 0.9524 | 0.8696 | 0.9252 |
| SVM with 41 genes | 0.9286 | 0.9091 | 0.9524 | 0.9302 | 0.9410 |
| SVM with 51 genes | 0.9286 | 0.9091 | 0.9524 | 0.9302 | 0.9410 |
| SVM with 148<br>genes | **0.9524** | 0.9130 | **1.0000** | **0.9545** | 0.9410 |
| SVM with 449<br>genes | **0.9524** | 0.9130 | **1.0000** | **0.9545** | 0.9138 |

Note: This table depicts the performance of various models on the test set. Within the listed models, TransGeneSelector was trained both in its unoptimized and optimized forms, unoptimized means without the implementation of 'early stop' optimization. Random Forest was trained with feature engineering on models using sets of 8, 11, 41, 51, 128, and 449 genes respectively. The SVM was trained utilizing genes chosen by the Random Forest model. 'AUC' stands for Area Under the Curve, indicating the probability that a randomly selected positive instance ranks higher than a randomly selected negative one. The highest values are highlighted in bold.

### TransGeneSelector demonstrates strong capability in key gene mining

In our study, the capability of TransGeneSelector’s Transformer network to mine key genes was thoroughly evaluated. We employed the SHAP method to analyze feature importance, highlighting the influence of each individual gene on the model’s predictions, and subsequently isolating the key genes involved in seed germination. Additionally, we utilized the Wrapper method in Random Forest to identify crucial genes in the germination process and compared the expression patterns of the identified genes by these two methods across all samples (Fig. 4). For Random Forest, we selected n_features_to_select parameters (gene numbers) that corresponded to cross-validation classification accuracies of 1.0, specifically 11, 51, 148, and 449, and contrasted them with the same number of genes chosen by TransGeneSelector’s SHAP method.

**Fig. 4.**
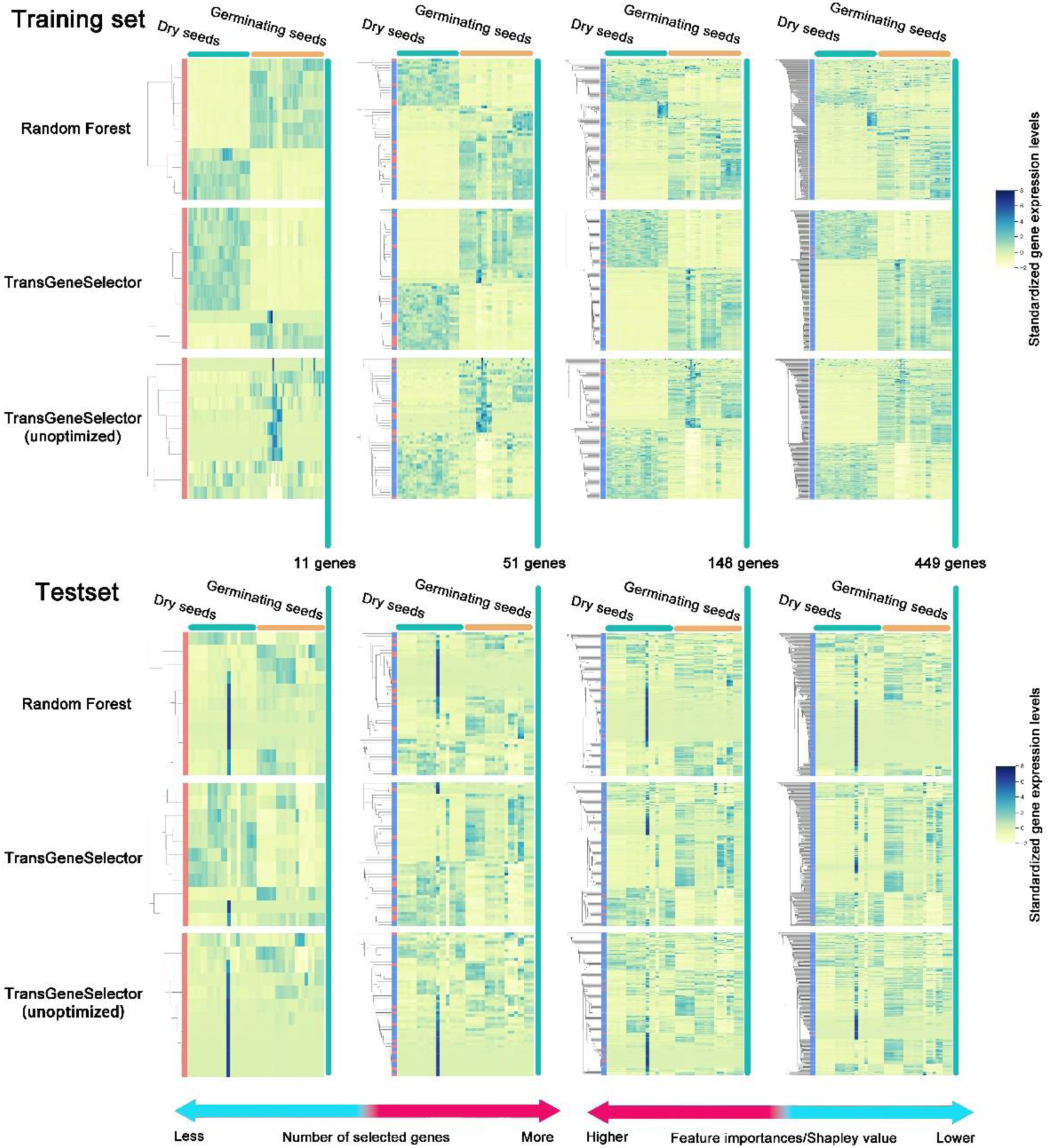
Comparative Expression Patterns of Genes Selected by TransGeneSelector and Random Forest. This heatmap illustrates the expression levels of genes selected by TransGeneSelector and Random Forest with varying numbers of genes. The color intensity represents the expression level, with darker shades indicating higher expression. TransGeneSelector’s selection is based on descending Shapley values, whereas Random Forest’ selection relies on different settings for the ‘n_features_to_select’ parameter. Unoptimized means TransGeneSelector was trained without the implementation of ‘early stop’ optimization.

The comparative analysis of expression patterns for training set (Fig. 4) yielded several noteworthy observations. Initially, when the number of genes targeted for selection was low, such as 11 and 51, the Random Forest’s chosen genes exhibited pronounced differences in expression patterns between dry seeds and germinating seeds. The expression levels within individual groups (either dry seeds or germinating seeds) were highly consistent, enabling clear differentiation from the other group (Fig. 4). This consistency was notably superior to that of genes selected by TransGeneSelector for the same gene numbers in the earlier examination. However, as the number of targeted genes increased to 148, the expression patterns of the genes identified by Random Forest began to appear disordered, and the distinctions between germinating and dry seeds became more subdued (Fig. 4). This trend became even more accentuated when the targeted genes reached 449 (Fig. 4).

Conversely, TransGeneSelector exhibited consistency in expression patterns for training set regardless of the number of targeted genes. Whether dealing with top 11, 51, 148, or 449 genes as designated by SHAP values, the expression patterns of genes selected by TransGeneSelector remained orderly and homogeneous within each group (dry seeds or germinating seeds), demonstrating evident distinctions between the groups (Fig. 4). This indicated a maintained uniformity and clear differentiation between germinating and dry seeds across varying gene numbers. The findings highlight a distinct divergence between TransGeneSelector and Random Forest in their characteristics for key gene mining. When expression patterns of the training set serve as the sole index for evaluating the efficacy of key gene selection, TransGeneSelector exhibits a clear advantage over Random Forest in the selection of genes across a range of targeted gene numbers.

We further evaluated the expression patterns based on the test set (Fig. 4). Contrary to the trends observed within the training set, the gene expression profiles, in general, did not exhibit as clear a distinction between the two groups on the test set. For Random Forest, regardless of whether a small or large number of genes were selected, the genes manifested a degree of inconsistency within groups. The expression profiles were more muddled, and clear demarcation between the groups became elusive. This lack of distinction was pervasive across varying gene quantities, indicating possible overfitting to the training data or the complex nature of the underlying gene interactions which Random Forest might not capture effectively in this context.

TransGeneSelector, while not achieving the same level of performance as observed in the training set, consistently showcased superior gene selection on the test set across all targeted gene numbers (Fig. 4). Genes chosen by TransGeneSelector not only maintained better internal group coherence but also succeeded in differentiating between the two groups, validating its robustness in gene mining and predictive capabilities.

An interesting phenomenon was observed in the context of the TransGeneSelector model that wasn’t optimally tuned (i.e., without early stopping for optimal model selection) (Fig. 4). For this unoptimized version, when selecting a smaller gene count, such as 11 genes, there was a tendency for these genes to exhibit pronounced expression in specific samples within the training set. This focused expression implies a possible over-reliance on certain training data patterns. Conversely, with a higher gene number, the expression profiles across groups in the training set remained distinct and orderly. However, when these genes were evaluated on the test set, their expression patterns became more chaotic, suggesting the potential pitfalls of operating with an unoptimized model. In conclusion, TransGeneSelector outshines Random Forest on both the training and test sets, underlining its superior ability to generalize across different data scenarios. To dissect the regulatory relationships among the 885 genes identified by both Random Forest and TransGeneSelector, we employed the MERLIN gene regulatory network construction algorithm (Fig. 5a). Intriguingly, when constructing the regulatory network using larger datasets unrelated to seed and seed germination, a majority (68%) of the regulatory relationships were observed from genes of Random Forest to those of TransGeneSelector (Fig. 5a and b). Yet, a pronounced shift materialized when we pivoted to using the specific dry seed and seed germination datasets from our study to inform the network construction. Not only did the number of regulatory relationships remain robust, but a reversal in transition patterns was evident. Specifically, 68% of the regulatory interactions were from genes of TransGeneSelector to those of Random Forest, starkly contrasting the 32% going the other way around (Fig. 5b). Further scrutiny revealed an intriguing detail: when leveraging the germination-related dataset, many of these regulatory relationships from TransGeneSelector to Random Forest were newly emergent, absent in the networks built from germination-unrelated datasets (Fig. 5a). This suggests that when the dataset resonates closely with germination processes, the constructed gene regulatory network is finely attuned to the nuances of seed germination relationships. Such findings underscore the prowess of TransGeneSelector. In the context of germination, it appears to effectively spotlight genes with heightened upstream regulatory influence, showcasing a marked advantage over Random Forest in discerning genes pivotal to seed germination dynamics.

**Fig. 5.**
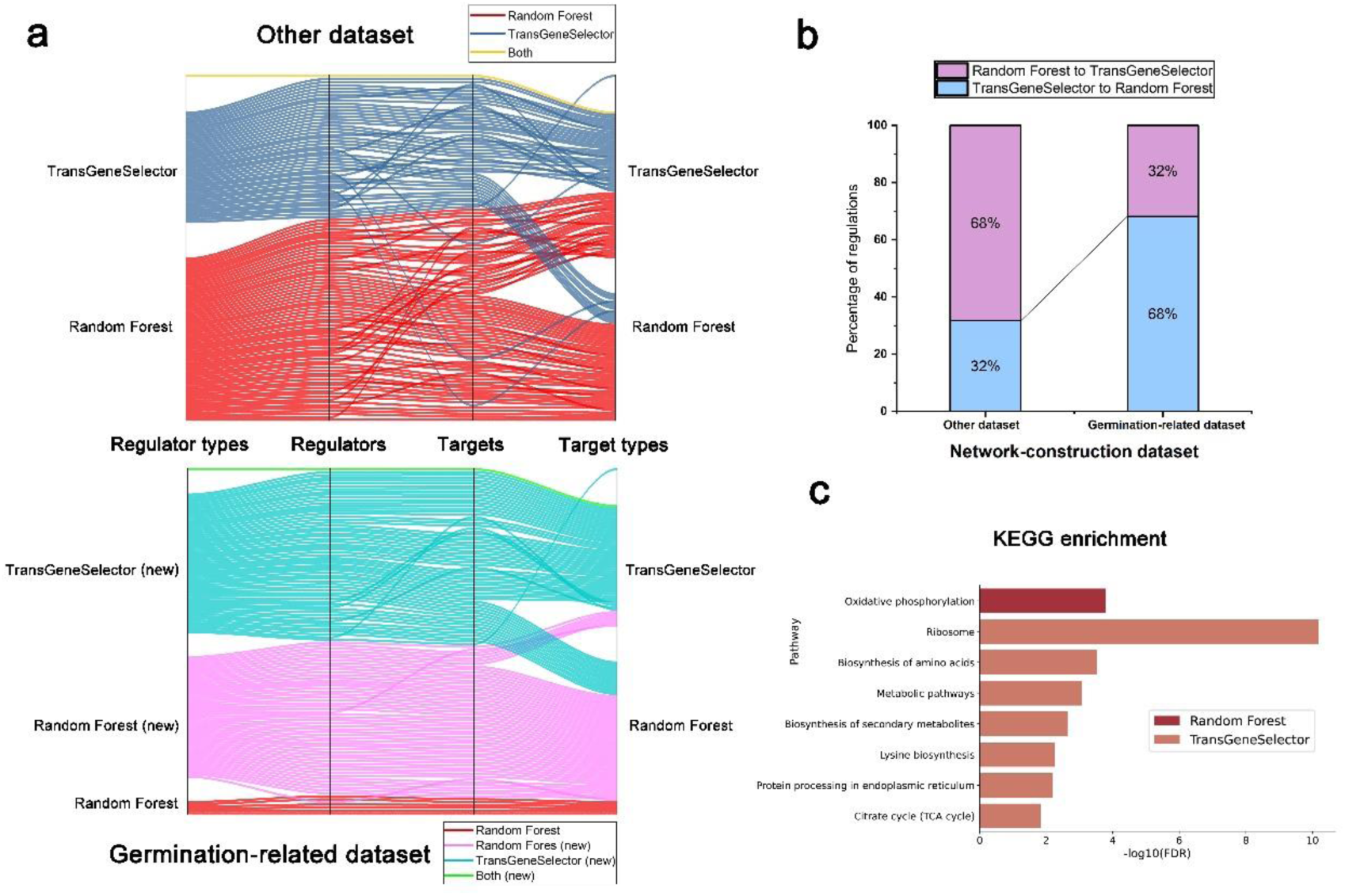
Comparative Analysis of Gene Regulatory Networks and Functional Enrichment for Genes Identified by TransGeneSelector and Random Forest. **a** Parallel Plot Illustrating Gene Regulatory Relationships: This segment showcases the MERLIN-derived regulatory links between genes identified by TransGeneSelector and Random Forest. Networks were constructed using datasets unrelated to germination (top) and those related to germination (bottom). Varied line colors signify regulatory relationships stemming from different types of regulators. Relationships labeled with (new) indicate newly emerged regulatory interactions identified within the germination-related dataset-constructed network. **b** Stacked Percentage Bar Chart of Regulatory Relationships Between Genes Identified by TransGeneSelector and Random Forest. **c** Comparative Results of KEGG Enrichment Analysis for Genes Identified by TransGeneSelector and Random Forest.

The Kyoto Encyclopedia of Genes and Genomes (KEGG) emphasizes the relationships between genes and metabolic pathways, rendering it an ideal platform for elucidating the interactions and functions of genes. In our research, we annotated and enriched each set of 449 genes mined by Random Forest and TransGeneSelector, focusing on these relationships (Fig. 5c). The KEGG enrichment results revealed that the genes extracted by Random Forest were only significantly concentrated in one pathway—Oxidative phosphorylation—while those extracted by TransGeneSelector were prominently enriched in seven pathways, namely, Ribosome, Biosynthesis of amino acids, Metabolic pathways, Biosynthesis of secondary metabolites, Lysine biosynthesis, Protein processing in endoplasmic reticulum, and Citrate cycle (TCA cycle) (Fig. 4c). The genes identified by both methods exhibited a blend of similarities and distinctions. Initially, both enriched pathways encapsulated aspects related to energy metabolism, such as Oxidative phosphorylation and Citrate cycle (TCA cycle). The genes mined by Random Forest, were exclusively enriched in the Oxidative phosphorylation pathway. This particular pathway is chiefly involved in energy metabolism and ATP synthesis and represents a downstream stage in the energy generation process, utilizing the products of other metabolic pathways like NADH and FADH2 to produce ATP(van Waveren and Moraes, 2008). Conversely, the genes unearthed by TransGeneSelector were enriched in the Citrate cycle (TCA cycle) pathway, which are key to central carbon metabolism processes, including glycolysis and the pentose phosphate pathway. This pathway generate precursor metabolites and reducing equivalents (NADH and FADH2) that feed into the Oxidative phosphorylation pathway(Tian et al., 2018). As such, it can be regarded as upstream of the Oxidative phosphorylation process.

In addition, genes significantly enriched in the pathways related to protein synthesis, such as Ribosome, Biosynthesis of amino acids, Lysine biosynthesis, and Protein processing in endoplasmic reticulum are of crucial importance for seed germination. During seed germination, which heralds the beginning of diverse plant life activities, a plethora of enzymes are required, along with various associated proteins to execute physiological functions transitioning from dormancy to an active state. Among these processes, protein synthesis relying on the mRNA stored within the seed is a crucial method to produce these key proteins(Marcus and Feeley, 1964; Fountain and Bewley, 1976; Shutov and Vaintraub, 1987; Müntz et al., 2001; Galland et al., 2014; Oracz and Stawska, 2016). Also noteworthy is the Biosynthesis of secondary metabolites pathway, which governs the synthesis of various secondary metabolites essential in numerous biological processes. This pathway signifies the onset of life diversity and bears a close relationship to seed germination(Reigosa and Pazos-Malvido, 2007; Čihák et al., 2017; Guerriero et al., 2018). Thus, in our research, genes identified by TransGeneSelector were notably enriched in several critical pathways associated with seed germination. In contrast, genes unearthed by Random Forest were predominantly concentrated in a more downstream pathway. This illustrates that TransGeneSelector is adept at pinpointing a broader and more upstream collection of genes crucial to seed germination than Random Forest, which is consistent with the gene regulatory network construction analysis.

### RT-qPCR test of genes identified by TransGeneSelector in response to germination

To elucidate the specific germination responses of genes identified by TransGeneSelector and Random Forest in *A. thaliana*, we performed RT-qPCR expression analyses (Fig. 6). Seeds were subjected to three germination conditions: dark germination, low-light germination, and high-light germination. For each condition, expression profiles were captured at four distinct germination time points: 0h (dry seeds), 12h, 24h, and 48h.

**Fig. 6.**
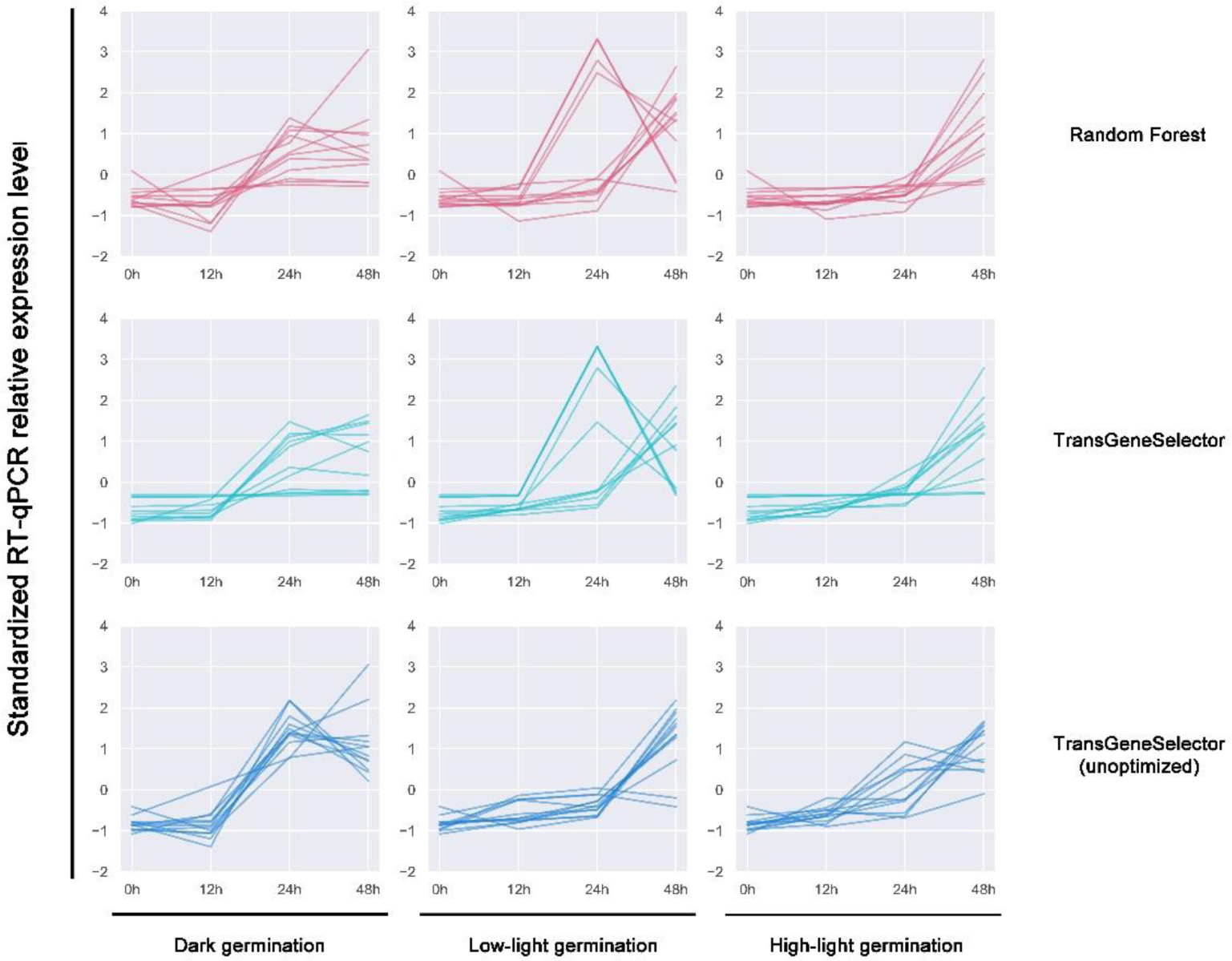
RT-qPCR quantitative analysis of the top 11 genes selected by TransGeneSelector and Random Forest in *A. thaliana* seeds under various germination conditions (n = 3) This figure presents the real-time quantitative polymerase chain reaction (RT-qPCR) results of the top 11 genes chosen by TransGeneSelector, Dark germination refers to seed germination conducted under full-black conditions, characterized by being wrapped in aluminum foil, simulating total darkness; Low-light germination refers to seed germination under a subdued illumination of 100 µmol photons of photosynthetic light, reflecting a low-light environment; High-light germination refers to seed germination under a more intense illumination of 200 µmol photons of photosynthetic light. Unoptimized means TransGeneSelector was trained without the implementation of ‘early stop’ optimization.

The results indicated that genes identified by both methods exhibited increased expression levels concurrent with the progression of germination, reinforcing the proficiency of both TransGeneSelector and Random Forest in capturing genes intimately associated with the germination process. A closer inspection of the data revealed that genes selected by the optimized TransGeneSelector exhibited expression patterns closely mirroring those chosen by Random Forest across all germination conditions.

Conversely, genes selected by the unoptimized TransGeneSelector exhibited distinct differences, most notably showing heightened expression at the 24h mark under dark germination conditions, which then declined (Fig. 6). Such a phenomenon was absent in genes identified by both the optimized TransGeneSelector and Random Forest (Fig. 6). Interestingly, the unoptimized TransGeneSelector demonstrated a tendency to selected genes with elevated expression specifically in the training set samples of *A. thaliana* seeds germinating under dark conditions (Supplementary Figure 1), these genes’ expression were consistent with genes related to dark germination of *A. thaliana* such as *KAI2* (Waters and Smith, 2013; Michael et al., 2022). This observation from the RT-qPCR result suggests that the unoptimized TransGeneSelector has a bias towards genes preferentially expressed during dark germination, as opposed to genes representative of a broader germination spectrum.

Furthermore, the expression patterns of genes identified by the optimized TransGeneSelector were characterized by higher consistency and uniformity than those selected by Random Forest. This observation further underscores the superior efficacy of TransGeneSelector over Random Forest in the domain of gene discovery related to seed germination.

Above all, while both methods have their merits, TransGeneSelector undoubtedly demonstrates enhanced potential than Random Forest, especially in terms of generalization and capturing universal germination-related gene expression dynamics. However, a note of caution remains on the importance of meticulous model optimization to fully harness its capabilities.

## Dicussion

In the realm of life sciences, particularly within plant research, gene mining holds an esteemed position. Historically, this endeavor begins with the analysis of limited sample data (Fregene et al., 2001; Wang et al., 2017; Westerman et al., 2022). This constraint has long presented challenges as conventional machine learning methods often overlook the complex mutual regulatory relationships between genes and omitted important genes, while existing deep learning techniques tend to struggle in these small sample size scenarios. To surmount these obstacles, our study introduces TransGeneSelector, the first deep learning method tailored for key gene mining in small transcriptomic datasets designed to effectively mine upstream key regulatory genes for specific biological processes within such constrained datasets. This innovation bridges the deficiencies of both traditional machine learning and deep learning methods in unearthing key genes from limited samples, and it has been successfully applied to uncover key upstream regulatory genes for seed germination in *A. thaliana*.

The development of TransGeneSelector started with a combination of data augmentation using the WGAN-GP network, quality control with an additional classifier, and the utilization of a simplified Transformer network structure. Impressively, we accomplished high-performance classification between dry and germinating seeds in a small dataset of 79 samples. After meticulous tuning of key hyperparameters like the number of augmented samples, it achieved superior accuracy, precision, and F1 scores compared to baseline Transformer models without generative augmentation. This highlights the benefits of strategically expanding limited training data through generative adversarial networks to enrich representation. Compared to conventional machine learning model of Random Forest and SVM, TransGeneSelector demonstrated competitive or better performance, validating its effectiveness for complex biological problems where acquiring abundant training data remains challenging.

Moreover, through the application of the Transformer network’s multi-head self-attention mechanism and the SHAP interpretable machine learning method, TransGeneSelector reliably identify pivotal genes related to the intricate process of seed germination, outperforming Random Forest. It consistently selected genes with orderly expression patterns that clearly distinguished dry and germinating states of seeds across varying targeted gene numbers. This stability was verified through test set evaluation, in contrast to Random Forest’s deteriorating performance, especially as gene count increased. KEGG pathway enrichment analysis revealed wider coverage of diverse upstream processes central to seed germination for genes selected by TransGeneSelector, whereas Random Forest focused on more downstream pathways. Furthermore, construction and analysis of the gene regulatory network using the MERLIN algorithm revealed intriguing insights. When using data unrelated to seed germination for network construction, most regulatory relationships were from Random Forest to TransGeneSelector genes. However, when employing the seed germination dataset, these relationships reversed, with most flowing from TransGeneSelector to Random Forest genes. Notably, many were new emergent relationships absent previously. This suggests TransGeneSelector better captures intricate regulatory dynamics specific to seed germination, and further proved that TransGeneSelector can reliably discover key upstream regulatory genes that exert major influence over downstream genes pivotal to seed germination.

RT-qPCR experiments validated that the genes identified by TransGeneSelector effectively captured key dynamics associated with seed germination. In contrast to higher variability in Random Forest selections, TransGeneSelector genes showed highly consistent and uniform expression changes responding to major germination conditions and progression. This further verifies TransGeneSelector’s balanced and robust gene discovery capabilities centered on the core biological process of interest. Despite these successes, TransGeneSelector, comprising a complex system of three separate neural networks, does present challenges in operational complexity and room for improvement. For instance, optimizing parameters in the WGAN-GP module and suitable threshold selection in the additional classifier requires refinement, as improper tuning could lead to biased results like the gene expression patterns observed in unoptimized models. This affirms the importance of meticulous optimization to avoid training data overfitting. Currently designed for binary classification tasks, future research must also explore multi-classification tasks, aligning with the needs of small-sized data applications.

In conclusion, the advent of TransGeneSelector in key gene mining within small sample transcriptomic data marks a significant advancement, offering a robust tool for mining key genes involved in vital life processes. By bridging the gaps of traditional methodologies and infusing the strengths of deep learning, this study not only furnishes a potent practical tool but also inaugurates a new perspective in gene mining research for plant and other organisms using Transformer networks.

## Materials and Methods

### TransGeneSelector framework

TransGeneSelector includes three neural networks, respectively, a sample generation network based on Wasserstein GAN with Gradient Penalty (WGAN-GP), an additional classifier network with a fully connected neural network architecture, and a classification network based on the Transformer architecture.

WGAN-GP, the sample generation network of TransGeneSelector, is an improvement over the original Wasserstein GAN (WGAN) (Goodfellow et al., 2014; Arjovsky et al., 2017) that addresses the limitations of the original model by using a gradient penalty instead of weight clipping to enforce the Lipschitz constraint. This results in more stable training and better convergence properties. For the original Wasserstein GAN, the loss function of the discriminator (critic)is defined as:

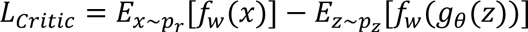

Where *f*_*w*_ is the discriminator (critic), *g*_*θ*_ is the generator, *p*_*r*_ is the real data distribution.

*p*_*z*_ is the noise distribution.

For the generator, the loss function is defined as:

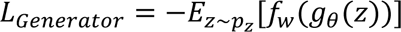

The WGAN model is trained by solving the following optimization problem:

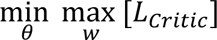

The WGAN-GP loss function is defined as:

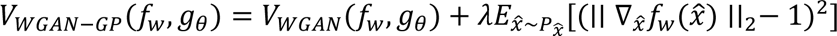

Where *V*_*WGAN*_(*f*_*w*_, *g*_*θ*_) is the original WGAN loss function, which is designed to address the limitations of the standard GAN loss function by using the Wasserstein distance instead of the Jensen-Shannon divergence. It consists of two parts: one for the discriminator (critic) and one for the generator. *f*_*w*_ is the discriminator (also called critic) in the WGAN-GP model. *g*_*θ*_ is the generator in the WGAN-GP model. *λ* is a hyperparameter that controls the strength of the gradient penalty term. 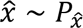 is the expectation over random samples 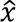 drawn from the distribution 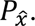 In WGAN-GP, 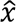 is a randomly weighted average between a real data point and a generated data point. 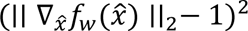 is the gradient penalty term. It penalizes the squared difference between the gradient norm of the discriminator with respect to its input 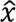 and the target norm value 1. The purpose of this term is to enforce the Lipschitz constraint on the discriminator, which helps to stabilize the training and improve convergence properties.

The WGAN-GP model is trained by solving the following optimization problem:

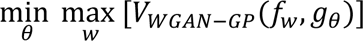

After generating the fake samples, the additional classifier network with a fully connected neural network architecture is used to filter out the fake samples and obtain high-quality samples. The network architecture consists of several fully connected layers (also known as linear layers) with Rectified Linear Unit (ReLU) activation functions in between, followed by a final linear layer with a Sigmoid activation function to output a probability value between 0 and 1. Here’s the mathematical representation of the additional classifier network:

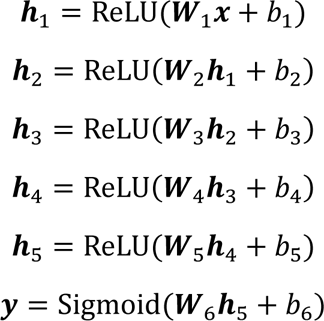

Where ***h***_*i*_ represents the output of the *i*^*th*^ hidden layer, ***W*** and *b* are the weight matrix cnd bids vector for the *i*^*th*^ layer, respectively. ReLU is the Rectified Linear Unit activation function, defined as:

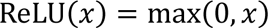

Sigmoid is the Sigmoid activation function, given an input *x*, the output *σ*(*x*) of the Sigmoid function is calculated as:

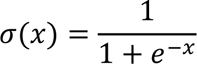

The network takes an input vector and passes it through the layers to produce a single output probability value. The output value can be thresholded to obtain the high-quality generated samples.

After the above sample-generating processes, the generated samples and real samples for seeds under both kinds of conditions (germinating or dry seeds) were altogether input into the Transformer network for biological process classification. It is start by using a fully connected network to reduce the dimensionality of the gene expression data for the numerous number of genes. The output of this step is a lower-dimensional representation of the input genes. Given an input expression value ***x***, the output ***y*** of a fully connected layer with weights ***W*** and biases *b* is calculated as

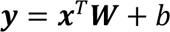

The lower-dimensional representation is then positional encoded to provide the Transformer network with information about the order of representation. The formula used for calculating the positional encoding values is as follows:

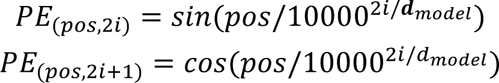

where *pos* is the position of the word in the sequence, *i* is the index of the dimension pair, and *d*_*model*_ is the dimension of the input embeddings.

When the lower-dimensional representation of the gene expression data is positional encoded, it is then fed into the Transformer Encoder. The Encoder processes the input sequence and produces a continuous representation, or embedding, of the input. The Transformer Encoder consists of multiple self-attention and feed-forward layers, allowing the model to process and understand the input sequence effectively. The multi-head self-attention mechanism in the encoder allows the model to attend to different parts of the input sequence simultaneously. It computes multiple attention outputs in parallel and then concatenates them before passing them through a linear transformation. The multi-head attention can be represented as:

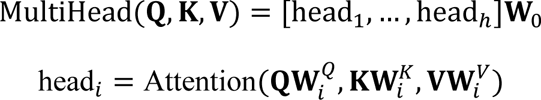

Here, **Q**, **K**, **V** represent the query, key, and value matrices, respectively, and **W** are the learnable parameter matrices. The scaled dot-product attention computes the attention scores by taking the dot product of the query and key matrices, dividing the result by the square root of the key vector dimension, and then applying a softmax function:

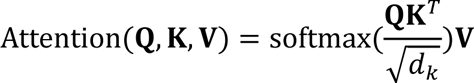

After the multi-head self-attention mechanism, the output is passed through a position-wise feed-forward network, which consists of two linear layers with a ReLU activation function in between. The position-wise feed-forward network can be represented as follows:

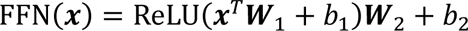

Where *x* is the input, *W*_1_and *W*_2_ are the weight matrices, and *b*_1_and *b*_2_ are the bias terms.

Finally, residual connections and layer normalization are applied after both the multi-head self-attention and position-wise feed-forward network to stabilize the training process and improve the model’s performance. Residual connections are used to allow gradients to flow through a network directly. The residual connection formula is:

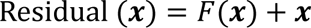

where *F*(***x***) is the output of the previous layer and ***x*** is the input

Layer normalization is applied to stabilize the training process. The layer normalization formula is:

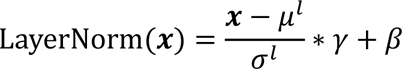

where *μ*^*l*^ and *σ*^*l*^ are the mean and standard deviation of the layer, respectively, and *γ* and *β* are learnable scale and shift parameters.

After processing the positional encoded lower-dimensional representation of the gene expression data through the Transformer Encoder, we use the first token of the output for classification, which is considered to contain the most relevant information for classification. We applied the Sigmoid function to the first token of the Encoder output, which is defined as:

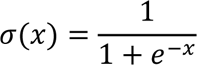

The Binary Cross-Entropy (BCE) loss function is then used as loss function for binary classification of dry seeds and germinating seed. It measures the dissimilarity between the predicted probability distribution and the true binary labels of a dataset. The BCE loss function is particularly useful when the output is a probability value between 0 and 1.

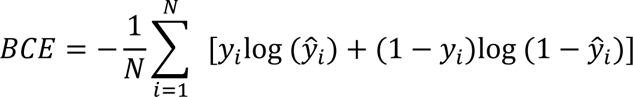

Where *N* is the number of samples in the dataset. *y*_*i*_ is the true binary label of the *i*^*th*^ sample (1 for the positive class and 0 for the negative class). 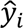 is the predicted probability of the *i*^*th*^sample belonging to the positive class.

To take into account the non-linearity of the activation functions and maintain the standard deviation of the activations around 1, we used the Kaiming Uniform Initialization (He et al., 2015) for initializing the weights of the Transformer network, the Kaiming Uniform Initialization initializes the weights from a uniform distribution *U*(−*a*, *a*), where:

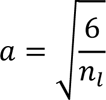

### Data description and processing

Data for our study were extracted from the NCBI GEO (Gene Expression Omnibus) database (https://www.ncbi.nlm.nih.gov/gds/) and the Expression Atlas database (https://www.ebi.ac.uk/gxa/experiments). For the training set of TransGeneSelector and Random Forest models, we selected experiments GSE116069, GSE161704, GSE163057, GSE167244, and GSE179008 from NCBI. These contained raw counts of 79 samples derived from dry seeds and germinating seeds, comprising 43 positive samples and 36 negative samples. The test set of these models were obtained from GSE167502, GSE230392, GSE94457, and GSE151223 with 42 samples, comprising 21 positive samples and 21 negative samples. The transcript-per-million (TPM) was calculated for each gene, utilizing its length and raw counts, and we retained only those genes present in all samples. Samples from the germinating group were designated as positive (1), and those from the non-germinating group were marked as negative (0). For additional MERLIN network analysis, we included experiments E-CURD-1, E-GEOD-30720, E-GEOD-52806, E-GEOD-64740, E-MTAB-4202, E-MTAB-7933, E-MTAB-7978 from Expression Atlas and GSE199116 from NCBI. These experiments included 268 samples unrelated to seed germination, and we proceeded with the same TPM calculations, retaining genes found across all samples.

### Benchmarking and evaluation metrics

To optimize the parameters of TransGeneSelector in the neural networks, a comprehensive grid search was performed on each part of the TransGeneSelector using the dry seed and germinating seed training set.

WGAN-GP: The epochs were set from 200 to 6000 in steps of 200, with combinations of learning rates (0.1, 0.01, 0.001). Model performance was assessed using the loss curve, the Fréchet Inception Distance (FID), and Uniform Manifold Approximation and Projection (UMAP) visualization to determine the best parameters.

Additional Classifier: Epochs (100, 150) and learning rate (0.1, 0.01) combinations were evaluated.

Transformer Network: Various combinations of embedding and header numbers (72/8, 240/8, 72/16, and 240/16), learning rates (0.1, 0.01, 0.001), and training periods (7, 21, and 35) were assessed. The best parameter combination was determined by the validation set loss values. To comprehensively compare the performance of optimized and unoptimized models, we trained TransGeneSelector with two additional model groups - one with early stopping criteria implemented for optimization, and one without early stopping, representing unoptimized models.

The Random Forest model parameters were optimized through training set using grid search to combine n_estimators (10, 100) with 200 values of n_features_to_select, uniformly spaced between 1 and 500. The best parameter combination was chosen based on model accuracy.

Both TransGeneSelector and Random Forest models were evaluated using a 5-fold cross-validation approach. For TransGeneSelector, a comprehensive assessment of performance was conducted using metrics including accuracy, precision, recall rate, and F1 score. In contrast, the evaluation of the Random Forest model focused solely on accuracy through cross-validation, with other metrics omitted as the Random Forest often achieved consistent 1.0 accuracy.

We utilized the test set data to evaluate the performance of all models by calculating the accuracy, precision, recall, F1 score, and AUC value. Through comparison of these metrics, the capabilities of different models were comprehensively assessed.

### Gene mining method

We applied SHAP (SHapley Additive exPlanations) (Lundberg and Lee, 2017) to mine important genes through Transformer network of TransGeneSelector by calculating the contribution of each gene to the prediction. SHAP is based on the concepts of game theory and can be applied to any machine learning model. The method uses Shapley values, which are derived from cooperative game theory, to fairly distribute the "payout" (i.e., the prediction) among the features. The formula for the Shapley value of gene *j* is given by:

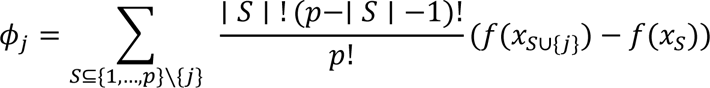

Here, *S* represents a subset of genes excluding gene *j*, *p* is the total number of genes, ∣ *S* ∣ is the number of genes of S, *S* ∪ {*j*} represents a new subset formed by adding gene *j* to the subset *S*, *f*(*x*) is the prediction function of the model. Those genes with high Shapley values were considered as important genes.

We performed Kyoto Encyclopedia of Genes and Genomes (KEGG) enrichment analysis on the genes mined from the TransGeneSelector and Random Forest, using the STRING v11.5 database (<http://www.string-db.org). We focused on pathways with FDR < 0.05, considering them as significantly enriched.

### Network Analysis

In this study, the Modular Regulatory Network Learning with Per Gene Information (MERLIN) algorithm(Roy et al., 2013) was employed to infer the regulatory network of genes mined using TransGeneSelector or Random Forest. The transcriptome data from the same dry seed and germinating seed datasets, as well as an additional dataset unrelated to seed or seed germination of *A. thaliana*, were prepared (as detailed in the Data Description and Processing section). Initially, the FPKM of the genes was transformed into Transcripts Per Million reads (TPM), and the mean expression level of each gene was calculated. Subsequently, the expression levels of the genes were zero-mean transformed. The MERLIN algorithm retained only those genes that (1) varied in expression value by at least ± 1 from the mean in at least five samples and (2) were included in the list of genes obtained in this study. All the genes were defined as both regulators and targets. A total of 10 sub-sets were created from the amended data matrix, with each sub-set containing 50% of the samples randomly selected from the complete matrix. Data from each sub-set were used to infer a MERLIN interaction. In the final MERLIN network, edges (which indicate the relationship between two genes) that appeared at least six times in the 10 sub-sets (confidence = 60%) were retained.

### Plant germination and RT-qPCR test

We used the same collection of *A. thaliana* col-0 mature seeds for both the germination experiment and the RT-qPCR quantitative analysis experiment. The seeds were stored at the Plant Development and Molecular Laboratory of Hunan Normal University, China.

We first selected surface-sterilized *A. thaliana* seeds, sown on 9-cm plates containing solidified 0.5 MS medium (pH 5.9), stratified in the dark at 4°C for 2 d, and then exposed to different light strengths and durations. The light strengths were set from weak to strong, corresponding to aluminum foil wrapped full-black germination conditions, 100 µmol photons photosynthetic light, and 200 µmol photons photosynthetic light. For germination time, we set 0h (dry seeds), 12h, 24h, and 48h. The temperature for germination was set to 25 ℃.

Seed samples were ground to a fine powder in liquid nitrogen. The ground tissue was transferred to a pre-chilled 1.5-mL Eppendorf tube, and total RNA was isolated using the TRIzol R Reagent (Life Technologies, Carlsbad, CA, United States), according to the manufacturer’s instructions. The RT-qPCR assay was conducted as described previously (Huang et al., 2020). Forward primers used for genes are listed in Supplementary Table 1, and actin1 was used as reference. The expected size of the amplified fragments varied from 80 to 200 bp. Three biological replicates were performed for each sample. Statistical analysis was performed using Piko Real Software 2.0. After standardization of each gene expression, the heatmap drawing was performed using Python.

## Data availability

The data utilized in this study were sourced from publicly available repositories. The gene expression data were extracted from the NCBI GEO (Gene Expression Omnibus) database (https://www.ncbi.nlm.nih.gov/gds/) and the Expression Atlas database (https://www.ebi.ac.uk/gxa/experiments). Specifically, the experiments GSE116069, GSE161704, GSE163057, GSE167244, and GSE179008 from NCBI were selected for the training set of the TransGeneSelector and Random Forest models, and the test set of these models were obtained from GSE167502, GSE230392, GSE94457, and GSE151223. For additional MERLIN network analysis, experiments E-CURD-1, E-GEOD-30720, E-GEOD-52806, E-GEOD-64740, E-MTAB-4202, E-MTAB-7933, E-MTAB-7978 from Expression Atlas and GSE199116 from NCBI were included. These datasets are publicly accessible and can be found in the respective databases.

## Acknowledgements and Fundings

The study was supported by the Natural Science Foundation of Hunan Province (2023JJ30436 and 2022JJ50249), Central Guidance Fund for Science and Technology Development in Hunan (2023ZYC012), Scientific Research Fund of Hunan Provincial Education Department (22A0487), Key Research Project of Hunan University of Arts and Science (E06022005)

We extend our gratitude to Li Zeng for his valuable advice and guidance on the paper.

## Author Contributions

X.J., Y.W., and K.H. conceived the basic idea and designed the framework. K.H. wrote the backbone code of TransGeneSelector. K.H., J.T., L.S., and P.X. carried out benchmark experiments. K.H. and J.T. carried out RT-qPCR test. S.Z., A.D., and P.M. prepared seed samples and performed seed germination test. Z.Z., M.J., and G.L. designed the figures. K.H. wrote the manuscript. All authors participated in the interpretation and writing of the manuscript.

## Materials & Correspondence

All requests for materials, data, and further information should be directed to, and will be fulfilled by, the corresponding author, Xiaocheng Jiang and Yun Wang. Inquiries regarding the specifics of the methodologies, datasets, or other related questions can be addressed to the same.

## Supplemental Materials

Supplementary Dataset 1

Supplementary Table 1 Primers of Genes Utilized for RT-qPCR Analysis.

Supplementary Figure 1 Transcriptomic Expression Patterns of the Top 11 Genes Identified by the Unoptimized TransGeneSelector in the Training Set

